# Combined Nanodrops Imaging and Ultrasound Localization Microscopy for Detecting Intracerebral Hemorrhage

**DOI:** 10.1101/2024.10.02.616087

**Authors:** Bing-Ze Lin, Alexander Changyu Fan, Yike Wang, Matthew R. Lowerison, Zhijie Dong, Qi You, Nathiya Vaithiyalingam Chandra Sekaran, Daniel Llano, Mark Borden, Pengfei Song

## Abstract

**Objective:** Advanced imaging methods are crucial for understanding stroke mechanisms and discovering effective treatments to reduce bleeding and enhance recovery. In preclinical *in vivo* stroke imaging, MRI, CT, and optical imaging are commonly used to evaluate stroke outcomes in rodent models. However, MRI and CT have limited spatial resolution for rodent brains, and optical imaging is hindered by limited imaging depth of penetration. Here we introduce a novel contrast-enhanced ultrasound imaging method to overcome these challenges and characterize intracerebral hemorrhage with unique insights.

**Methods:** We combined microbubble (MB)-based ultrasound localization microscopy (ULM) and nanodrop (ND)- based vessel leakage imaging to achieve simultaneous microvascular imaging and hemorrhage detection. ULM maps brain-wide cerebral vasculature with high spatial resolution and identifies microvascular impairments around hemorrhagic areas. NDs are sub-micron liquid-core particles which can extravasate due to blood-brain barrier (BBB) breakdown, serving as positive contrast agents to detect hemorrhage sites.

**Results:** Our findings demonstrate that NDs could effectively accumulate in the hemorrhagic site and reveal the location of the bleeding areas upon activation by focused ultrasound beams. ULM further reveals the microvascular damage manifested in the form of reduced vascularity and decreased blood flow velocity across areas affected by the hemorrhagic stroke.

**Conclusion:** The results demonstrate that sequential ULM combined with ND imaging is a useful imaging tool for basic *in vivo* research in stroke with rodent models where brain-wide detection of active bleeding and microvascular impairment are essential.

## Introduction

Intracerebral hemorrhage (ICH) represents more than 10% of the 17 million occurrences of stroke globally each year. It is associated with a mortality rate exceeding 40%, and only 20% of the survivors achieve functional independence within six months ^1,2^. ICH stands as the second most frequent and most lethal form of stroke, yet effective medically or surgically proven treatments remain elusive ^1^. Due to blood vessel rupture and bleeding into the brain parenchyma, hemorrhages usually lead to an increasing cerebral pressure and subsequent hypoperfusion ^3^. Early detection of alterations in cerebral blood flow (CBF) and microcirculation can provide valuable insights into the underlying mechanisms of hemorrhage. Impairments in microcirculation can result in altered microvascular perfusion and tissue oxygenation, potentially leading to tissue damage and neurological disorders. While CT and MRI are powerful imaging modalities for human and large animal studies, their spatial resolution is insufficient for capturing the microvascular range, making direct measurement of microvascular density and hemodynamics unfeasible^4–7^. As such, there is a strong preclinical need for whole-brain high-resolution imaging techniques that can provide detailed insights into blood flow and the dynamics of hemorrhage at the microvascular level.

Ultrasound imaging offers an enticing solution for this clinical need because it is widely available in both clinical and non-clinical settings. Ultrasound (e.g., transcranial Doppler) is regularly utilized in the clinic for characterizing major cerebral vessels^8^ but conventional techniques lack sufficient spatial resolution for detecting small vessel abnormalities and also lack extravascular imaging capabilities for detecting vessel leakage.

Ultrasound localization microscopy (ULM) is an emerging super-resolution ultrasound imaging technique that utilizes the localization and tracking of intravascular-injected MBs to image microvessels in deep tissues^9,10^. In addition to structural imaging of microvessels, ULM also leverages MB tracking to measure blood flow velocity^11^. Previous work by Chavignon *et al*. used ULM to characterize ischemic and hemorrhagic stroke in a rat model ^12^ and demonstrated the effectiveness of using ULM for detecting microvascular perfusion losses during acute onset of stroke. ULM was also used in humans for stroke imaging by Demene *et al*., who demonstrated the feasibility of detecting deeply seated aneurysms in stroke patients with transcranial ULM^13^. One common limitation associated with these pioneering works is the use of MBs as contrast agent for stroke imaging^12,14^. Contrast MBs are micron-sized gas-core spheres that are too large to extravasate to provide vessel leakage measurements commonly associated with hemorrhagic stroke. Although quantitative microvascular diffusion indices may be derived to differentiate hemorrhagic vs. ischemic stroke, they require prolonged imaging and therefore do not facilitate early detection. As such, a method that offers both direct assessment of vessel leakage and microvascular impairment associated with hemorrhagic stroke remains elusive.

Phase-changing nanodrops (NDs) offer a promising solution for detecting active bleeding in hemorrhagic stroke due to their smaller size and longer circulation time than MBs^15^. Their reduced size allows them to evade filtration by macrophages in both the cardiopulmonary circulation (e.g., lungs), and the systemic circulation (e.g., liver, and spleen), enhancing their effectiveness in prolonged circulation^16–19^. NDs were previously used for ULM applications to enhance microvasculature imaging performance due to their smaller size, which facilitates their perfusion into smaller vessels ^20,21^. Upon acoustic activation, non-echogenic NDs expand into echogenic MBs to provide adequate backscattering for detection ^22,23^. NDs have also been used in cancer applications to provide vessel leakage measurements, as well as serving as a vehicle for drug delivery ^24 25^. Inspired by the existing work and the unique features of NDs, here we propose to combine ND-based vessel leakage imaging with ULM to provide direct and comprehensive evaluation of microvascular impairment associated with stroke. Unlike MBs, NDs have the capability to extravasate from the vasculature. Their small size and perfluorocarbon liquid core facilitate a long circulation half-life, allowing ample time for NDs to extravasate. In addition, owing to their small size, a higher ND count can be delivered than MBs at the same particle volume, which effectively enhances the concentration of NDs in the blood stream to increase overall extravasation. These advantages make NDs ideal candidates for vessel leakage detection in the context of hemorrhagic stroke.

In this study, the efficacy of ND imaging was initially assessed in a tissue-mimicking flow phantom. A hemorrhagic stroke mouse model based on collagenase injection was subsequently developed to evaluate the feasibility of using NDs for detecting early-stage active bleeding. Concurrent ULM imaging was used to monitor alterations in microvascular density and flow velocity post hemorrhagic stroke. The combined utility of ULM and ND imaging shows great potential for improving the understanding of stroke pathology through detailed examinations of vascular integrity and blood leakage dynamics.

## Methods

### Phospholipid NDs preparation

NDs were fabricated at the University of Colorado Boulder using a previously described MB-condensation method^26^. In brief, lipid films were prepared by dissolving DSPC and DSPE-PEG2K (Avanti Polar Lipids) in a 9:1 molar ratio in chloroform (Sigma-Aldrich), followed by solvent evaporation under nitrogen gas. The films were dried overnight under vacuum and stored at −20°C. Next, the lipid films were rehydrated with 1X PBS to a lipid concentration of 2mg/mL, heated to 55°C, and sonicated in a water bath at 45°C until transparent. The suspension was cooled to 4°C and sonicated with a Branson 450 Digital Sonifier (Branson) under a perfluorobutane (FluoroMed) hood to generate MBs. The MBs were size-isolated by centrifugation to remove those with diameters greater than 3 µm and stored in crimp-sealed 2-mL serum vials^27^. To form droplets, the MB suspension was rapidly cooled in 70% isopropyl alcohol solution (Sigma-Aldrich) at −20°C for 2–3 minutes with periodic mixing. Vials were then pressurized with room-temperature perfluorobutane until the suspension turned transparent, indicating successful gas core condensation. The resulting NDs were transferred to syringes and maintained at 4°C during shipping to UIUC. ND size was assessed using NanoSight NS300 (Malvern) and Coulter Mutisizer 4e (Beckman Coulter).

### Flow phantom experiments

ND activation was verified using a flow phantom with channel’s diameter approximately 450 μm. Ultrasound imaging was conducted using a Verasonics Vantage 256 System (Verasonics) equipped with an L22-14vX, 128-element high-frequency linear transducer, operated at a center frequency of 15.625 MHz (Verasonics). Images were acquired using plane-wave compounding with steering angles ranging from -5 to 5 degrees in 1-degree increments, achieving a post-compounding frame rate of 1,000 Hz. The ND concentration was diluted to 5×10^8^ particles/mL and introduced into the flow phantom with a 1-mL syringe.

### Animal preparations

All animal procedures followed the University of Illinois IACUC guidelines (Protocol #22033). Twelve-week-old female C57BL/6 mice were used for the hemorrhagic model and ND imaging. Mice were anesthetized with 4% isoflurane in 100% oxygen for induction, followed by 1-2% for maintenance. The head was secured in a stereotaxic frame (Kopf), and body temperature was maintained at 37 °C with a heating pad (Physitemp). The scalp was removed, and temporalis muscles were dissected. A bilateral craniotomy window was created using a high-speed drill (Foredom). Artificial cerebrospinal fluid (aCSF) was applied to keep the brain tissue moist^28^.

### Hemorrhagic stroke mouse model establishment

An automatic nanoinjector (Nanoject II, Drummond) was mounted on a stereotaxic frame via a micromanipulator. A capillary glass needle (Borosilicate, 1.2 x 0.69 mm x 10 cm, Sutter) filled with mineral oil (Sigma-Aldrich) was inserted into the nanoinjector (Fig. 1a). Following this setup, a collagenase solution (0.075 U/0.4 µl in saline) (C0773, Sigma-Aldrich) was drawn into the needle, ensuring no air bubbles. Collagenase, commonly used to establish ICH rodent models^29,30^, was injected after craniotomy. The needle was aligned with the bregma, moved 2 mm left and 2 mm caudally, and injected with 100 nL at 1.2 mm depth or 25 nL at 0.6 mm depth. Once both the tail vein needle and the capillary glass needle were in position, the ND infusion commenced for 2 min, followed by the collagenase injection. The needle remained in place for 5 min to ensure collagenase deposition (Fig. 1a).

**Figure 1.**
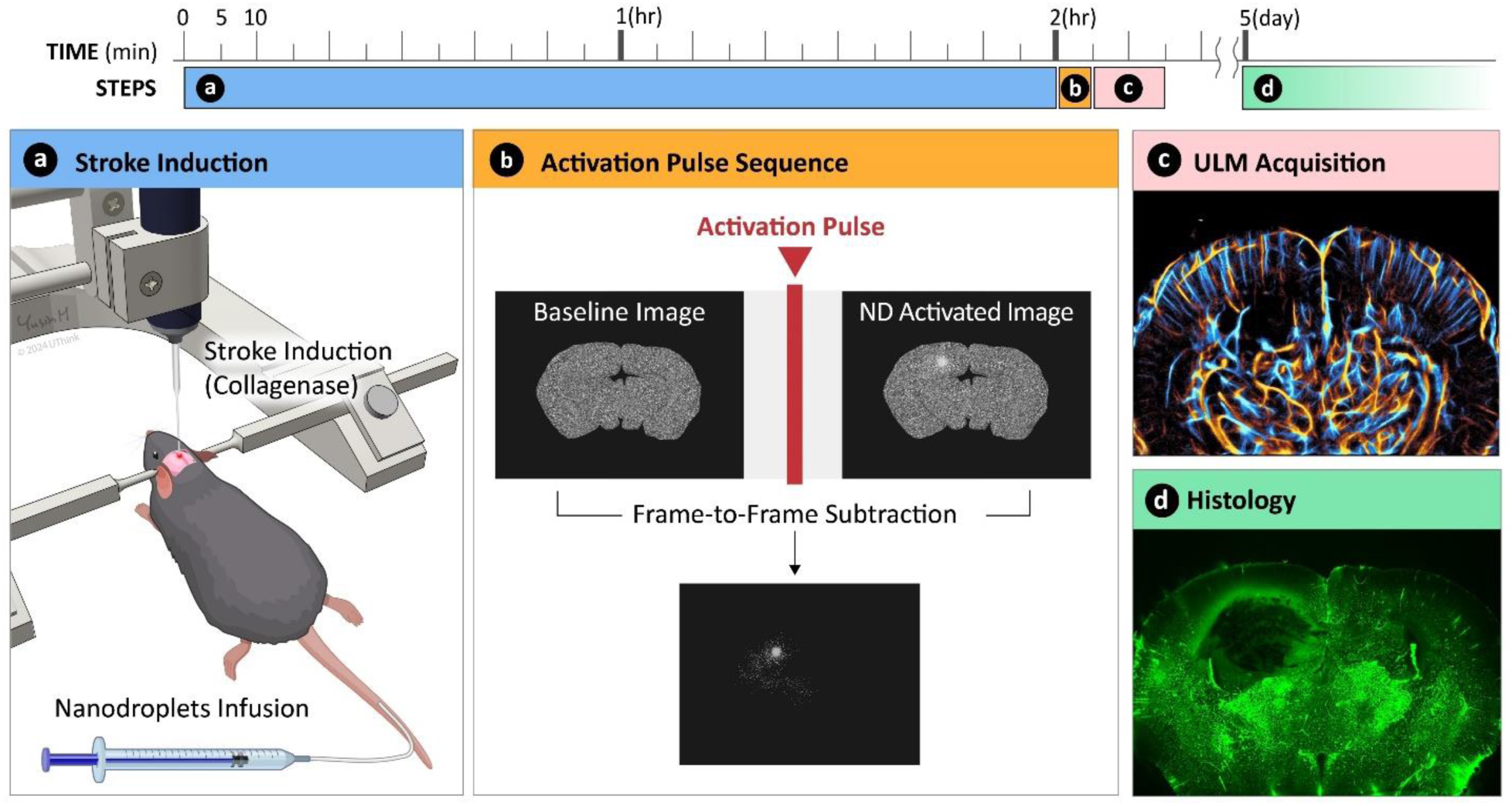
Schematic overview of stroke Induction and ultrasound-activated nanodrop imaging in a mouse Model. (a) Experimental timeline beginning with the induction of stroke through intracerebral collagenase injection and simultaneous intravenous infusion of NDs. (b) Application of activation pulses to induce nanodrop vaporization. (c) ULM imaging for cerebral vasculature. (d) Histological evaluation of the hemorrhagic region.

### ND activation sequence

NDs were administered via a 29-gauge tail vein catheter with a syringe pump (NE-300, New Era) infusing 1 mL of NDs at 200 µL/min (Fig. 1a). The mouse remained under anesthesia for 2 hours post-infusion to allow ND extravasation and clearance from blood vessels. ND imaging involved a special sequence: low-voltage (6V) plane wave for baseline imaging, high-voltage (30V) sweeping focused beams for ND activation (40 cycles at 4 mm focal depth, Fig. 2a), and another low-voltage sequence for post-activation imaging (Fig. 1b). The derated mechanical index (MI_0.5_) (0.5 dB/cm/MHz) of the center focused beam was measured at 0.54.

**Figure 2.**
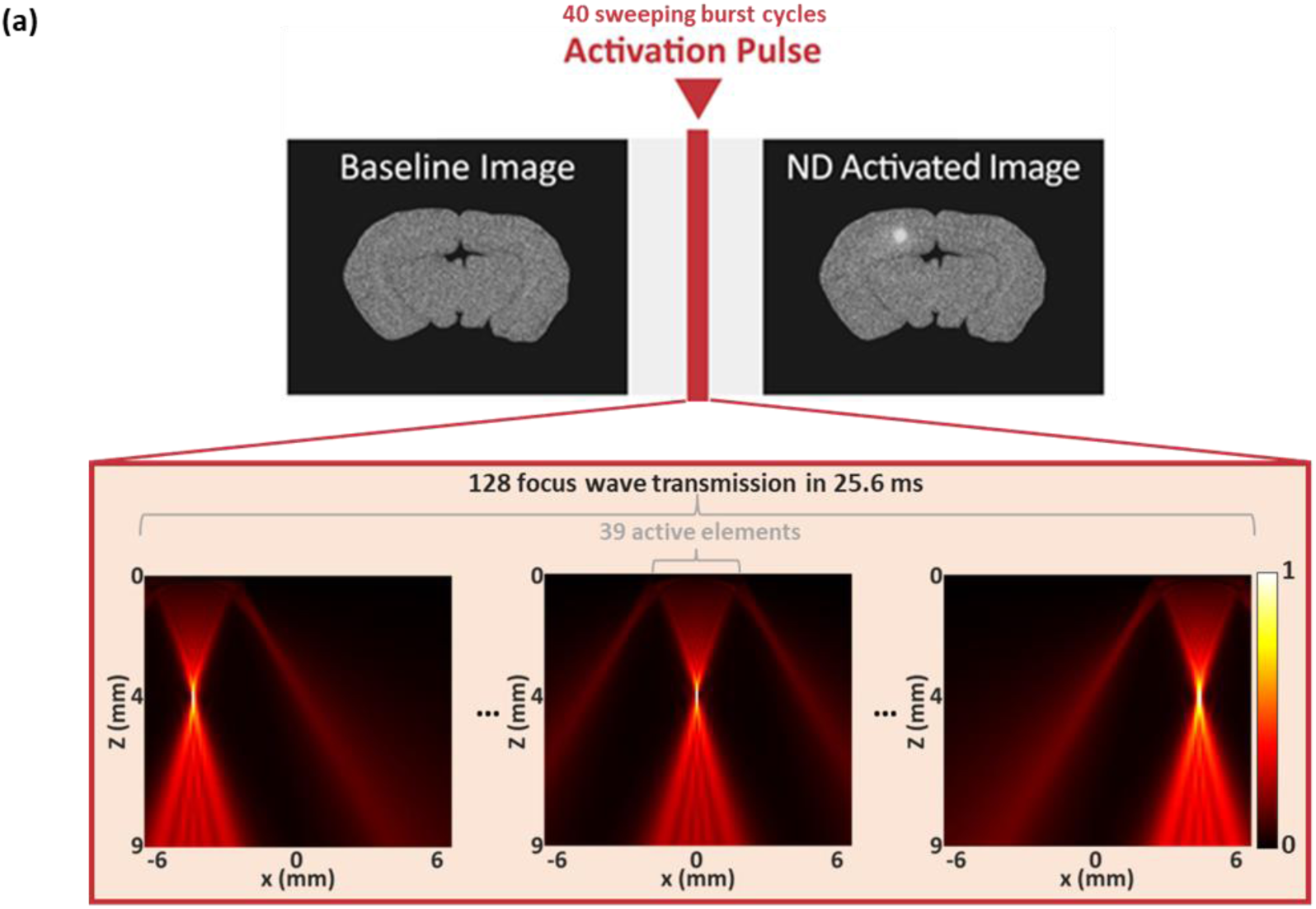
Schematics of the NDs imaging workflow. (a) Verasonics simulation was used to visualize the 128 focused beams used for ND activation.

The ND activation beam’s acoustic field was assessed through simulations using the Verasonics simulator and hydrophone measurements with the ONDA AIMS III system (Onda). The measurements employed a capsule hydrophone (HGL-0200, Onda) in degassed deionized water. The MI0.5 at the activation beam’s spatial peak was measured.

### ND detection algorithm

Each ND ultrasound image dataset contained 400 frames of baseline images and 400 frames of post-activation images (N=400). To isolate changes due to ND activation from baseline images, we performed frame-to-frame subtraction of the image at each pixel (x, y) to the first frame.

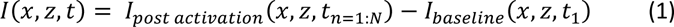

Equation (1) represents the difference between the post-activation frame and the baseline frame.

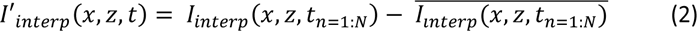

where *I*_*interp*_ is the spatially interpolated version of *I* (interpolation factor = 4), and *I*′_*interp*_ is the mean subtracted version of *I*_*interp*_ (Eq. 2).

To further improve spatial resolution and reduce uncorrelated spatiotemporal noise, the second-order cross-cumulant (C _second-order_) as introduced in SOFI^31,32^ was performed (Equation 3), which calculates the spatial coherence of adjacent pixels to preserve correlated signals (i.e., stationary extravasated NDs) and reject noise with low spatial coherence. As shown in Eq. 3, a local window (3 x 3 pixels) was used for the spatial coherence measurements between pixels with 400 temporal samples, and the results were averaged temporally across these samples.

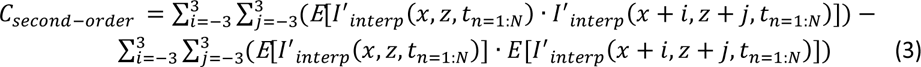

### ULM imaging acquisition and reconstruction

MBs (DEFINITY®, Lantheus) were introduced into the bloodstream through the tail vein catheter (Fig. 1c). A 5 mL MB syringe was attached to the pump, controlling infusion at 20 µL/min. After removing the glass needle, ultrasound gel was applied to the transducer, positioned over the collagenase injection site for scanning. Ultrasound data were collected with the same L22-14vX transducer using the ultrafast imaging sequence from the phantom study.

To separate MB signals from the surrounding tissue in each IQ (in-phase and quadrature) data, SVD clutter filtering was applied with an adaptively set singular value threshold ^33,34^. MBs moving towards and away from the transducer were separated into two datasets for directional information using a directional filter^35^. The filtered MB data were cubic-interpolated to achieve a resolution with 9.86 µm pixel size, approximately one-tenth of the wavelength at 15 MHz. Each interpolated frame underwent 2D normalized cross-correlation with the empirically determined point spread function (PSF) of the imaging system. MB center positions were identified as regional maxima in the cross-correlation results. MB centers were tracked across frames using the uTrack algorithm^36^, with a persistence control of 10 frames to ensure tracking accuracy. Microvessel density and flow velocity maps were then derived from the MB tracks for final ULM imaging results.

### FITC-Dextran Perfusion and Histology Imaging

After ND and ULM imaging, 200 µL of 10 mg/mL FITC-dextran (70,000Da, Sigma-Aldrich) in saline was administered via tail vein catheter. Mice were euthanized 3 minutes later, and brains were harvested, fixed in 10% formalin for 24 hours, and cryoprotected in 10%, 20%, and 30% sucrose (S0389, Sigma-Aldrich). Brains were sectioned at a thickness of 40 µm using a cryostat (CM1850, Leica). Sections were imaged with an Olympus IX71 fluorescence microscope (Redmond) using 2.5X and 10X lenses with excitation/emission wavelengths of 472 ± 30 nm/520 ± 35 nm (Fig. 1d). Region of interest (ROI) selections and quantitative analysis were performed using MATLAB (2020a).

## Results

### Analyzing ND Characteristics in a Flow Phantom

To evaluate the effectiveness of the ND activation sequence across the imaging region of interest (ROI), we examined the ND activation events in a tissue-mimicking flow phantom. The MB concentration was (7.82 ± 0.09) ×10^7^ particles/mL, and the mean MB diameter was 2.64 ± 0.01 µm (Fig. 3a). The ND concentration was (9.70 ± 0.54) ×10^8^ particles/mL, and the mean size of NDs was 134.3 ± 7.0 nm (Fig. 3b). Compared to MBs, NDs were significantly smaller in size, facilitating their extravasation through leaky vessels. Figs. 3c-j present the ND activation results with axial and lateral imaging views. Figs. 3c-d and g-h show that B-mode imaging does not provide the necessary contrast to visualize the activated NDs, which were clearly revealed using the subtraction method (i.e., Eqs. 1-3) as demonstrated in Figs. 3e-f and i-j. A single activation sequence consisting of 40 cycles of sweeping focused beams was used in this experiment. These results demonstrate the feasibility of using the ND imaging sequence and post-processing method for detecting NDs in a similar imaging depth as our *in vivo* study settings.

**Figure 3.**
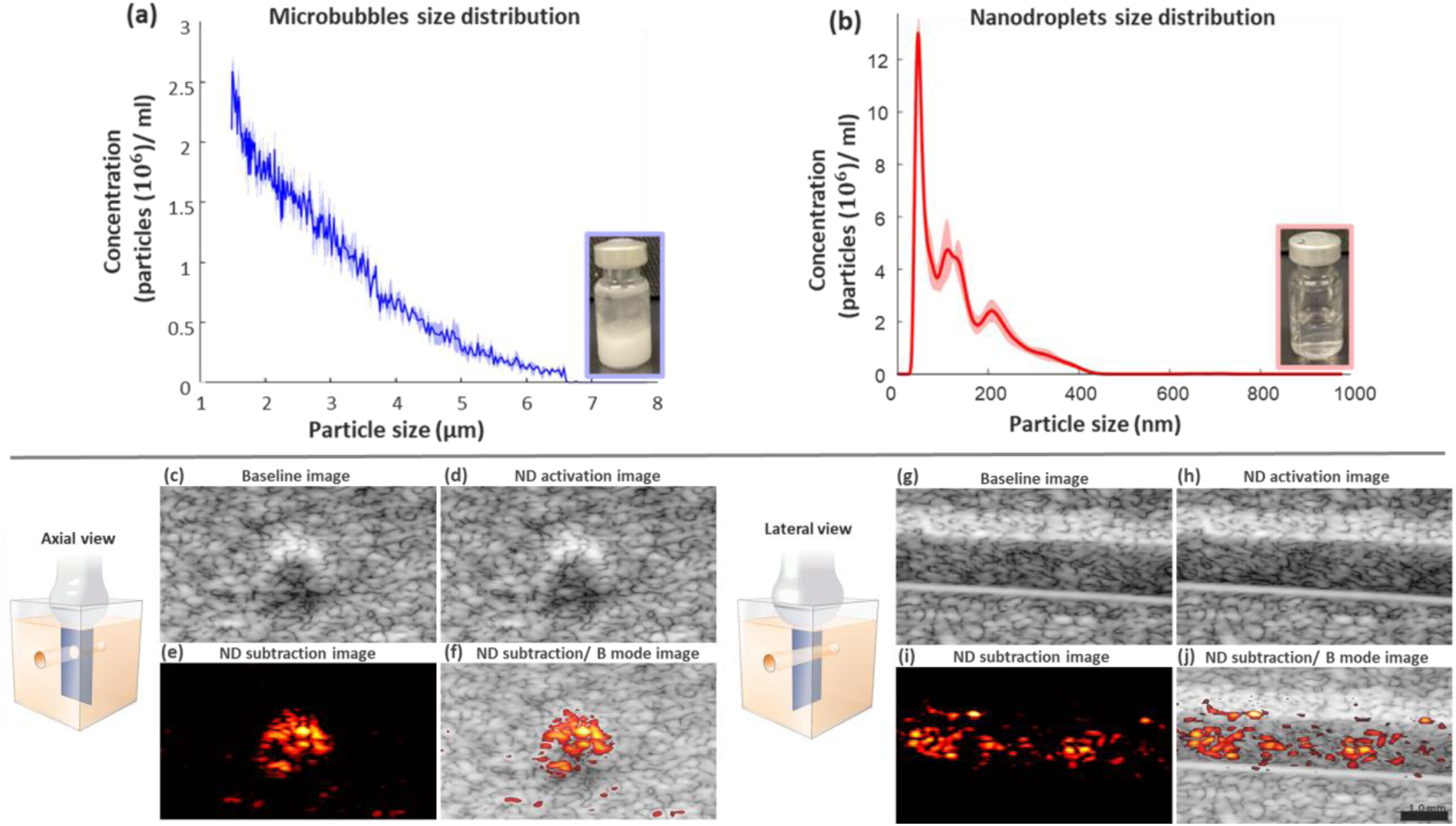
ND and MB size characterizations and flow phantom studies. (a) and (b) show the particle size distribution measurements for MBs and NDs. (c)-(j) show the ultrasound imaging results of NDs pre- and post-activation. Two different views (axial and lateral) were used for imaging.

### ULM Combined with ND Imaging for Hemorrhage Detection *In Vivo*

Fig. 4 shows the *in vivo* imaging results for the collagenase-induced hemorrhagic stroke model. The first row of Fig. 4 displays the results for the 100 nL collagenase injection, and the second row displays the results for the 25 nL injection. As can be clearly seen from both Fig. 4a and Fig. 4d by ULM imaging, hypoperfusion was successfully induced by collagenase in the left cortical region of the mouse brain (highlighted by the dashed boxes). The vascular damage is confirmed by histological imaging presented in Figs. 4c and f, which clearly demonstrate hyper-enhancement in the hemorrhagic area, indicating BBB breakdown and vessel leakage induced by collagenase and elevated leakage of FITC-dextran (Figs. 4c,f). Fig. 4c (100 nl collagenase injection) shows a larger hemorrhagic area with low fluorescence signals in the center region, which contains blood clots that did not effectively take up FITC-dextran. Collectively, these results indicate profound vascular damage induced by collagenase, which successfully induced hemorrhagic stroke in our study. Figs. 4b and e present the ND imaging results, from which it can be clearly seen that NDs successfully extravasated out of the leaky vessels in the stroke region. No significant ND signal enhancement was detected in normal brain tissues. Figs. 4b, c also demonstrate that the amount of ND extravasation is correlated to the dose of collagenase injection (quantitative analysis provided below), indicating a positive correlation between ND signal intensity and the severity of stroke.

**Figure 4.**
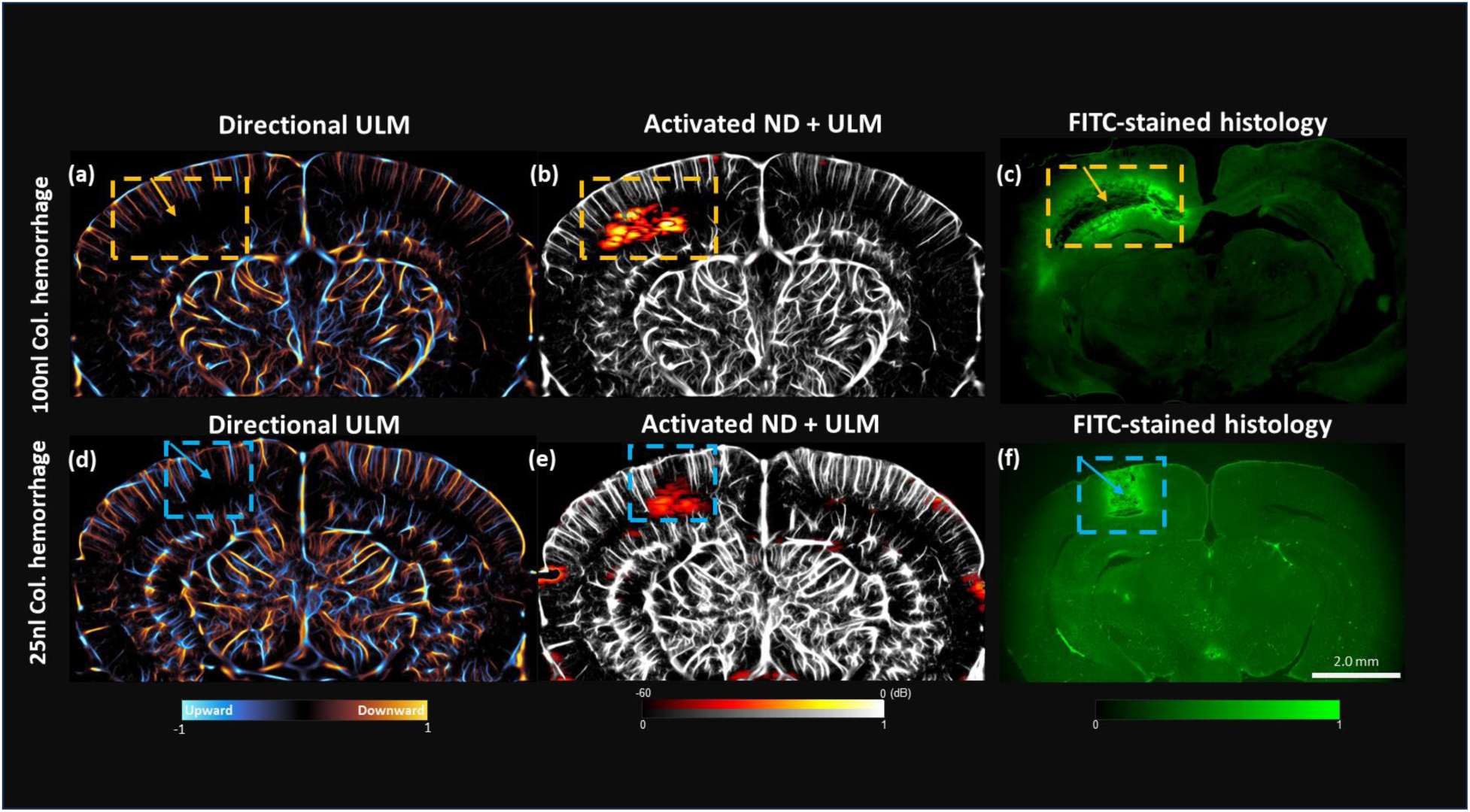
ULM and ND imaging in Mouse Intracranial Hemorrhage Models. (a, d) Directional ULM images showcase the cerebral vasculature and flow directions for both 100 nL and 25 nL collagenase-induced hemorrhagic models. The dashed boxes indicate the hemorrhagic stroke region. (b, e) show the ND imaging results (hot color map) overlaid on ULM images (identical to a and d but displayed with gray scale). (c, f) Fluorescein Isothiocyanate (FITC)-stained histological imaging results. The hemorrhagic sites were confirmed with hyper-enhancement which indicates leaky vessels.

### Quantitative Analysis of ND Imaging Results

To further quantify the ND imaging results, we superimposed the ND images over the corresponding ULM vascular and FITC-dextran histology images (Figs. 5a, b). Enlarged views highlighting the hemorrhagic regions are displayed in Figs. 5c, d (100 nL in central column, and 25 nL in right column). Figs. 5c,d clearly demonstrate the strong spatial correlations among ND extravasation, vascular deficiencies indicated by the hypoperfusion region in ULM, and vascular damages indicated by histology. Regions-of-interests (ROIs) were manually selected for the ND activation area (Figs. 5e, f) as well as the hyper-enhancement fluorescence area (Figs. 5g, h) for quantitative analysis. For the 100 nL and 25 nL collagenase injections, the area of FITC-dextran leakage is 2.69 mm² and 0.98 mm², respectively. Here we use the FITC-dextran leakage area as a quantitative measure of vascular leakage, which serves as the ground truth for ultrasound measurements. The ND extravasation area for the 100 nL and 25 nL injections is 1.77 mm² and 0.74 mm², respectively. The relative vessel leakage area measured by ND imaging between 100 nL and 25 nL (i.e., 1.77 vs. 0.74, which is appromiately 2.4-fold) is similar to that measured by FITC-dextran histology (i.e., 2.69 vs. 0.98, which is approximately 2.74-fold). In addition, the Jaccard similarity between the FITC-dextran histology image and ND extravasation image was evaluated, which produces a coefficient of 0.73±0.02 of the 100 nL case, and 0.72±0.04 for the 25 nL case. This result indicates good agreement between ND imaging and histology in the context of identifying the hemorrhagic stroke region with active bleeding.

**Figure 5.**
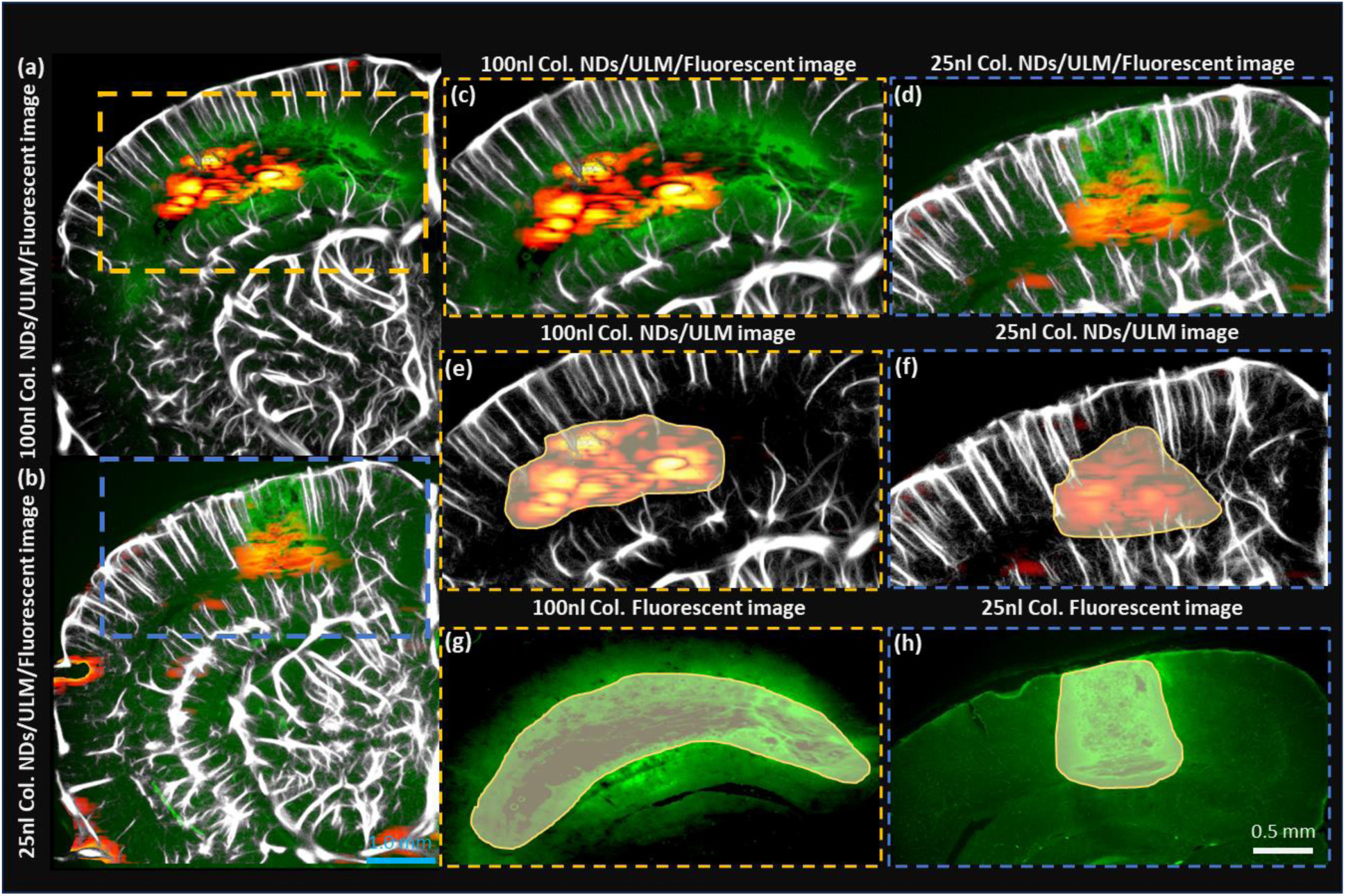
Regional analysis of ND imaging results in the mouse stroke model. (a, b) ULM images with ND and fluorescent imaging overlays for 100 nL (a) and 25 nL (b) collagenase injection, (c-d) present the enlarged views of the hemorrhagic region for the two collagenase dosages. (e-f) display the ND activation map overlaid on the ULM image, with ND area manually selected for quantitative analysis. (g-h) display the fluorescent images with the stroke region manually segmented for quantitative analysis.

### Quantitative Hemodynamic Analysis based on ULM Imaging

Variations in regional blood flow velocity and vascular density were further quantitatively analyzed from ULM imaging results. Blood flow velocity maps were generated by MB tracking and are presented in Fig. 6. In addition to the structural deficiencies, there is a clear reduction in blood flow velocity in the hemorrhage region compared to the control side (e.g., the regions identified by the red and green dashed boxes in Fig. 6a, b). For the 100 nL collagenase injection (Fig. 6e), the hemorrhagic side had an average flow velocity of 4.76 mm/s compared to 5.2 mm/s on the control side (Fig. 6g). Similarly, for the 25 nL injection (Fig. 6f), the hemorrhagic side averaged 5.40 mm/s versus 5.62 mm/s on the control side (Fig. 6h). Fig. 6i illustrates the differences in blood flow velocity between the control and hemorrhagic sides. In the 100 nL sample, the average blood flow velocity in the hemorrhagic side is significantly lower than that in the control side (p < 0.001). Similarly, in the 25 nL sample, the average blood flow velocity in the hemorrhagic side is also lower than that in the control side (p < 0.01). This finding suggests that larger hemorrhage volume led to a greater reduction in blood flow velocity. In addition to differences in flow velocity, we also observed differences in vessel density between the control and hemorrhagic sides. For the 100 nL injection, the vessel density on the hemorrhagic side was 16.87%, compared to 26.45% on the control side (a reduction of 9.58%), indicating a decreasing trend in vessel density (Fig. 6j). Likewise, for the 25 nL injection, the vessel density on the hemorrhagic side was 36.57%, while it was 42.33% on the control side (a reduction of 5.76%), also showing a comparable reduction. These results consistently demonstrate that the 100 nL model leads to a greater decrease in vessel density than the 25 nL model. These quantitative measurements indicate both structural (i.e., vessel density) and functional (i.e., blood flow speed) impairments to the cerebral vasculature due to the collagenase-induced hemorrhagic stroke.

**Figure 6.**
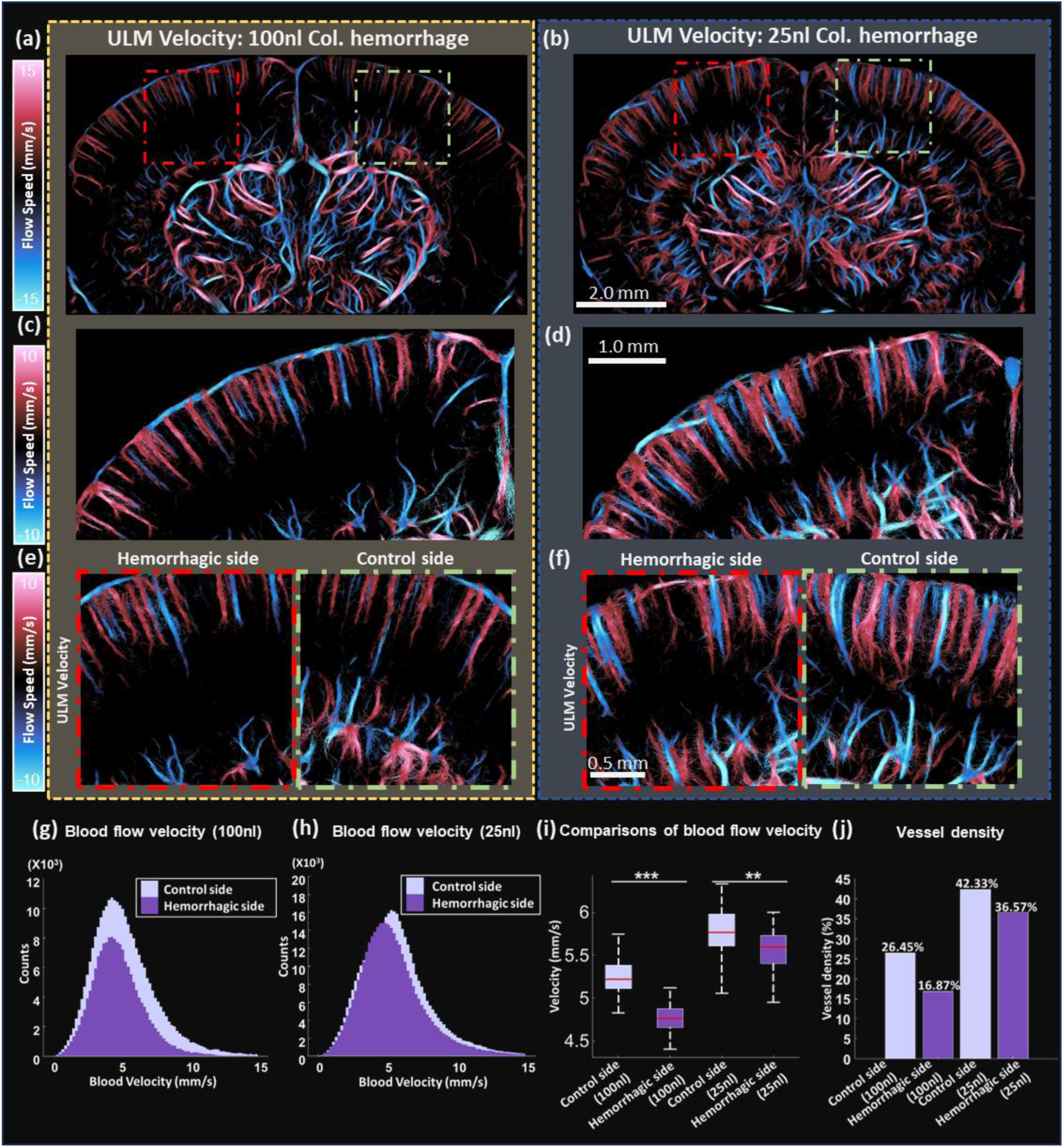
Hemodynamic assessment in the mouse hemorrhage model. (a, b) ULM blood flow velocity maps highlighting areas of normal flow (green box) and compromised flow (red box) induced by hemorrhage. (c, d) Magnified images showing flow speed within the hemorrhagic regions for 100 nL (c) and 25 nL (d) injections. (e, f) Magnified images of hemorrhagic (red box) and control regions (green box) for quantitative analysis in (g, h, i, j). (g, h) Blood flow velocity distribution for regions with vasculature from (e) 100 nL and (f) 25 nL, respectively. (i) Averaged blood flow velocity comparison between the control and hemorrhagic sides for regions with vasculature (e) 100 nL and (f) 25 nL injections, respectively. Significance is denoted as follows: *p < 0.05, **p < 0.01, ***p < 0.001. (j) Vessel density comparisons between the control and hemorrhagic sides for (e) 100 nL and (f) 25 nL injections.

## Discussion

In this study, we introduce a novel method for detecting hemorrhagic stroke using activated extravasated nanodrops (NDs) and ULM imaging in mice. Intracerebral hemorrhage was induced by cerebral injections of collagenase and detected with an intravenous infusion of NDs. An ND activation sequence with sweeping focused ultrasound beams was developed together with an ND detection algorithm based on signal subtraction and correlation. Fluorescence-based histology imaging was used as the ground truth to confirm hemorrhage. ULM was combined with ND extravasation imaging to provide evaluation of hemorrhage-induced hemodynamic impairments in the mouse brain.

Our results indicate that hemorrhagic stroke induced active leakage of NDs out of the vascular space, and the amount of ND extravasation was proportional to the dose of the collagenase. Hemorrhagic stroke also significantly undermined the structure and function of the cerebral vasculature, as indicated by the reduced microvessel density and blood flow speed. Similar to ND, the level of impairment to the cerebral vasculature was also proportional to the dose of the collagenase. These results indicate that the method introduced here has the potential to assess the severity and extent of hemorrhagic stroke.

ULM revealed acute and extensive vascular damage in the vicinity of the hematoma after the onset of the hemorrhagic stroke. The significant reduction in blood flow revealed by ULM can be explained by the physiological responses to hemorrhage, which involves the coagulation cascade and subsequent thrombin generation to stop bleeding^37,38^. Another cause of the blood flow reduction involves physical displacement or compression of brain tissues from the accumulating blood, which elevates the extravascular pressure and reduces blood flow^39^. Similar findings of reduction in blood flow have also been reported in existing literature^40^.

Our study has several limitations. Firstly, the timing of ND administration may impact the outcome of leakage detection. In this study, ND was allowed 2 h to extravasate before activation, which was sufficient for our study design. However, the dynamics of ND distribution could vary significantly with the extent of the BBB disruption and other physiological responses to the hemorrhage. These variabilities may impact the accuracy and sensitivity of hemorrhage detection using the proposed method. A future study is necessary to systematically study the impact of the timing of ND administration and imaging for different stroke characteristics.

Second, the bioeffect of ND vaporization remains unclear in this study. While our study did not observe adverse effects directly attributable to ND activation, the long-term effects of ultrasound-induced ND vaporization in brain tissues remain unexplored^41–43^. The potential for thermal or mechanical damage to surrounding tissues, especially with repeated or prolonged exposure, necessitates thorough investigation in future studies.

## Conclusions

Our study presents a novel method for detecting intracerebral hemorrhagic stroke by combining nanodrop (ND) imaging with ultrasound localization microscopy (ULM) in a mouse model. Our approach utilizes the ultrasound-activated NDs that extravasated and accumulated at the hemorrhage site for hemorrhage detection. In addition, ULM provided complementary insights of cerebrovascular structural and functional damages induced by hemorrhagic stroke. The proposed method has the potential to become a useful tool for differentiating hemorrhagic versus ischemic stroke as well as assessing the severity and extent of the hemorrhagic stroke.

## Acknowledgements

This work was supported in part by the National Institute of Biomedical Imaging and Bioengineering under Grant R21EB030072 and Grant R21EB030072-01S1, in part by the National Institute of Neurological Disorders and Stroke under Grant R56 NS131516, in part by the National Institute on Aging under Grant R21AG077173, and in part of Chan Zuckerberg Initiative (CZI) Ben Barres Early Career Acceleration Award. We thank Dr. Xinzhu Yu for assistance with surgical medicine preparation and acknowledge Yu-Sin Huang for illustrational contributions.

## Data availability Statement

The data can be made available upon request to Bing-Ze Lin.

## References

1. Broderick JP, Grotta JC, Naidech AM, Steiner T, Sprigg N, Toyoda K, et al. The Story of Intracerebral Hemorrhage. Stroke. 2021 May;52(5):1905–14.

2. Tsao CW, Aday AW, Almarzooq ZI, Anderson CAM, Arora P, Avery CL, et al. Heart Disease and Stroke Statistics-2023 Update: A Report From the American Heart Association. Circulation. 2023 Feb 21;147(8):e93–621.

3. Unnithan AKA, M Das J, Mehta P. Hemorrhagic Stroke. In: StatPearls [Internet]. Treasure Island (FL): StatPearls Publishing; 2024 [cited 2024 Feb 20]. Available from: http://www.ncbi.nlm.nih.gov/books/NBK559173/

4. Mannheim JG, Schlichthaerle T, Kuebler L, Quintanilla-Martinez L, Kohlhofer U, Kneilling M, et al. Comparison of small animal CT contrast agents. Contrast Media Mol Imaging. 2016 Jul;11(4):272–84.

5. Mannheim JG, Kara F, Doorduin J, Fuchs K, Reischl G, Liang S, et al. Standardization of Small Animal Imaging—Current Status and Future Prospects. Mol Imaging Biol. 2018 Oct 1;20(5):716–31.

6. Amirrashedi M, Zaidi H, Ay MR. Towards quantitative small-animal imaging on hybrid PET/CT and PET/MRI systems. Clin Transl Imaging. 2020 Aug 1;8(4):243–63.

7. Callewaert B, Jones EAV, Himmelreich U, Gsell W. Non-Invasive Evaluation of Cerebral Microvasculature Using Pre-Clinical MRI: Principles, Advantages and Limitations. Diagnostics. 2021 Jun;11(6):926.

8. Kirsch JD, Mathur M, Johnson MH, Gowthaman G, Scoutt LM. Advances in Transcranial Doppler US: Imaging Ahead. RadioGraphics. 2013 Jan;33(1):E1–14.

9. Errico C, Pierre J, Pezet S, Desailly Y, Lenkei Z, Couture O, et al. Ultrafast ultrasound localization microscopy for deep super-resolution vascular imaging. Nature. 2015 Nov;527(7579):499–502.

10. Christensen-Jeffries K, Browning RJ, Tang MX, Dunsby C, Eckersley RJ. In Vivo Acoustic Super-Resolution and Super-Resolved Velocity Mapping Using Microbubbles. IEEE Trans Med Imaging. 2015 Feb;34(2):433–40.

11. Lowerison MR, Huang C, Lucien F, Chen S, Song P. Ultrasound localization microscopy of renal tumor xenografts in chicken embryo is correlated to hypoxia. Sci Rep. 2020 Feb 12;10(1):2478.

12. Chavignon A, Hingot V, Orset C, Vivien D, Couture O. 3D transcranial ultrasound localization microscopy for discrimination between ischemic and hemorrhagic stroke in early phase. Sci Rep. 2022 Aug 26;12(1):14607.

13. Demené C, Robin J, Dizeux A, Heiles B, Pernot M, Tanter M, et al. Transcranial ultrafast ultrasound localization microscopy of brain vasculature in patients. Nat Biomed Eng. 2021 Mar;5(3):219–28.

14. Hingot V, Brodin C, Lebrun F, Heiles B, Chagnot A, Yetim M, et al. Early Ultrafast Ultrasound Imaging of Cerebral Perfusion correlates with Ischemic Stroke outcomes and responses to treatment in Mice. Theranostics. 2020 Jun 12;10(17):7480–91.

15. Borden MA, Shakya G, Upadhyay A, Song KH. Acoustic nanodrops for biomedical applications. Curr Opin Colloid Interface Sci. 2020 Dec 1;50:101383.

16. Sirsi SR, Borden MA. Microbubble compositions, properties and biomedical applications. Bubble Sci Eng Technol. 2009 Nov 1;1(1–2):3–17.

17. Zhang M, Fabiilli ML, Haworth KJ, Padilla F, Swanson SD, Kripfgans OD, et al. Acoustic Droplet Vaporization for Enhancement of Thermal Ablation by High Intensity Focused Ultrasound. Acad Radiol. 2011 Sep 1;18(9):1123–32.

18. Dove JD, Mountford PA, Murray TW, Borden MA. Engineering optically triggered droplets for photoacoustic imaging and therapy. Biomed Opt Express. 2014 Dec 1;5(12):4417–27.

19. Kulle A, Thanabalasuriar A, Cohen TS, Szydlowska M. Resident macrophages of the lung and liver: The guardians of our tissues. Front Immunol. 2022 Nov 30;13:1029085.

20. Zhang G, Harput S, Lin S, Christensen-Jeffries K, Leow CH, Brown J, et al. Acoustic wave sparsely activated localization microscopy (AWSALM): Super-resolution ultrasound imaging using acoustic activation and deactivation of nanodroplets. Appl Phys Lett. 2018 Jul 2;113(1):014101.

21. Riemer K, Tan Q, Morse S, Bau L, Toulemonde M, Yan J, et al. 3D Acoustic Wave Sparsely Activated Localization Microscopy With Phase Change Contrast Agents. Invest Radiol. 2023 Sep 11;10.1097/RLI.0000000000001033.

22. Moyer LC, Timbie KF, Sheeran PS, Price RJ, Miller GW, Dayton PA. High-intensity focused ultrasound ablation enhancement in vivo via phase-shift nanodroplets compared to microbubbles. J Ther Ultrasound. 2015;3:7.

23. Zhang P, Porter T. An *in vitro* Study of a Phase-Shift Nanoemulsion: A Potential Nucleation Agent for Bubble-Enhanced HIFU Tumor Ablation. Ultrasound Med Biol. 2010 Nov 1;36(11):1856–66.

24. Ho YJ, Yeh CK. Concurrent anti-vascular therapy and chemotherapy in solid tumors using drug-loaded acoustic nanodroplet vaporization. Acta Biomater. 2017 Feb 1;49:472–85.

25. Mannaris C, Bau L, Grundy M, Gray M, Lea-Banks H, Seth A, et al. Microbubbles, Nanodroplets and Gas-Stabilizing Solid Particles for Ultrasound-Mediated Extravasation of Unencapsulated Drugs: An Exposure Parameter Optimization Study. Ultrasound Med Biol. 2019 Apr 1;45(4):954–67.

26. Brambila CJ, Lux J, Mattrey RF, Boyd D, Borden MA, de Gracia Lux C. Bubble Inflation Using Phase-Change Perfluorocarbon Nanodroplets as a Strategy for Enhanced Ultrasound Imaging and Therapy. Langmuir. 2020 Mar 24;36(11):2954–65.

27. Feshitan JA, Chen CC, Kwan JJ, Borden MA. Microbubble size isolation by differential centrifugation. J Colloid Interface Sci. 2009 Jan 15;329(2):316–24.

28. Yu X, Nagai J, Marti-Solano M, Soto JS, Coppola G, Babu MM, et al. Context-Specific Striatal Astrocyte Molecular Responses Are Phenotypically Exploitable. Neuron. 2020 Dec;108(6):1146–1162.e10.

29. MacLellan CL, Silasi G, Poon CC, Edmundson CL, Buist R, Peeling J, et al. Intracerebral Hemorrhage Models in Rat: Comparing Collagenase to Blood Infusion. J Cereb Blood Flow Metab. 2008 Mar 1;28(3):516–25.

30. Krafft PR, Rolland WB, Duris K, Lekic T, Campbell A, Tang J, et al. Modeling Intracerebral Hemorrhage in Mice: Injection of Autologous Blood or Bacterial Collagenase. J Vis Exp JoVE. 2012 Sep 22;(67):4289.

31. Dertinger T, Colyer R, Vogel R, Enderlein J, Weiss S. Achieving increased resolution and more pixels with Superresolution Optical Fluctuation Imaging (SOFI). Opt Express. 2010 Aug 30;18(18):18875–85.

32. Dertinger T, Colyer R, Iyer G, Weiss S, Enderlein J. Fast, background-free, 3D super-resolution optical fluctuation imaging (SOFI). Proc Natl Acad Sci. 2009 Dec 29;106(52):22287–92.

33. Lowerison MR, Huang C, Kim Y, Lucien F, Chen S, Song P. In Vivo Confocal Imaging of Fluorescently Labeled Microbubbles: Implications for Ultrasound Localization Microscopy. IEEE Trans Ultrason Ferroelectr Freq Control. 2020 Sep;67(9):1811–9.

34. Song P, Manduca A, Trzasko JD, Chen S. Ultrasound Small Vessel Imaging With Block-Wise Adaptive Local Clutter Filtering. IEEE Trans Med Imaging. 2017 Jan;36(1):251–62.

35. Huang C, Lowerison MR, Trzasko JD, Manduca A, Bresler Y, Tang S, et al. Short Acquisition Time Super-Resolution Ultrasound Microvessel Imaging via Microbubble Separation. Sci Rep. 2020 Apr 7;10(1):6007.

36. Jaqaman K, Loerke D, Mettlen M, Kuwata H, Grinstein S, Schmid SL, et al. Robust single-particle tracking in live-cell time-lapse sequences. Nat Methods. 2008 Aug;5(8):695–702.

37. Iannucci J, Renehan W, Grammas P. Thrombin, a Mediator of Coagulation, Inflammation, and Neurotoxicity at the Neurovascular Interface: Implications for Alzheimer’s Disease. Front Neurosci [Internet]. 2020 Jul 24 [cited 2024 Mar 29];14. Available from: https://www.frontiersin.org/journals/neuroscience/articles/10.3389/fnins.2020.00762/full

38. Zhang R, Bai Q, Liu Y, Zhang Y, Sheng Z, Xue M, et al. Intracerebral hemorrhage in translational research. Brain Hemorrhages. 2020 Mar 1;1(1):13–8.

39. Kellner CP, Chartrain AG, Nistal DA, Scaggiante J, Hom D, Ghatan S, et al. The Stereotactic Intracerebral Hemorrhage Underwater Blood Aspiration (SCUBA) technique for minimally invasive endoscopic intracerebral hemorrhage evacuation. J NeuroInterventional Surg. 2018 Aug;10(8):771–6.

40. Yang GY, Betz AL, Chenevert TL, Brunberg JA, Hoff JT. Experimental intracerebral hemorrhage: relationship between brain edema, blood flow, and blood-brain barrier permeability in rats. J Neurosurg. 1994 Jul 1;81(1):93–102.

41. O’Brien WD. Ultrasound-biophysics mechanisms. Prog Biophys Mol Biol. 2007;93(1–3):212–55.

42. Vlatakis S, Zhang W, Thomas S, Cressey P, Moldovan AC, Metzger H, et al. Effect of Phase-Change Nanodroplets and Ultrasound on Blood–Brain Barrier Permeability In Vitro. Pharmaceutics. 2024 Jan;16(1):51.

43. Kee ALY, Teo BM. Biomedical applications of acoustically responsive phase shift nanodroplets: Current status and future directions. Ultrason Sonochem. 2019 Sep 1;56:37–45.

